# Interspecies variation in hominid gut microbiota controls host gene regulation

**DOI:** 10.1101/2020.08.17.255059

**Authors:** Amanda L. Muehlbauer, Allison L. Richards, Adnan Alazizi, Michael Burns, Andres Gomez, Jonathan B. Clayton, Klara Petrzelkova, Camilla Cascardo, Justyna Resztak, Xiaoquan Wen, Roger Pique-Regi, Francesca Luca, Ran Blekhman

**Affiliations:** Department of Genetics, Cell Biology and Development, University of Minnesota, Minneapolis, Minnesota, USA; Department of Ecology, Evolution and Behavior, University of Minnesota, Minneapolis, Minnesota, USA; Center for Molecular Medicine and Genetics, Wayne State University, Detroit, Michigan 48201, USA; Department of Biology, Loyola University, Chicago, Illinois, 60660, USA; Department of Animal Science, University of Minnesota, Saint Paul, Minnesota, USA; Department of Biology, University of Nebraska at Omaha, Omaha, Nebraska, USA; Department of Food Science and Technology, University of Nebraska-Lincoln, Lincoln, Nebraska, USA; The Czech Academy of Sciences, Institute of Vertebrate Biology, Brno, Czech Republic; Liberec Zoo, Liberec, Czech Republic; The Czech Academy of Sciences, Institute of Parasitology, Ceske Budejovice, Czech Republic; Department of Biostatistics, University of Michigan, Ann Arbor, MI, USA; Department of Obstetrics and Gynecology, Wayne State University, Detroit, Michigan 48201, USA

## Abstract

The gut microbiome exhibits extreme compositional variation between hominid hosts. However, it is unclear how this variation impacts host physiology, and whether this effect can be mediated through microbial regulation of host gene expression in interacting epithelial cells. Here, we characterized the transcriptional response of colonic epithelial cells *in vitro* to live microbial communities extracted from humans, chimpanzees, gorillas, and orangutans. We found most host genes exhibit a conserved response, whereby they respond similarly to the four hominid microbiomes, while some genes respond only to microbiomes from specific host species. Genes that exhibit such a divergent response are associated with relevant intestinal diseases in humans, such as inflammatory bowel disease and Crohn’s disease. Lastly, we found that inflammation-associated microbial species regulate the expression of host genes previously associated with inflammatory bowel disease, suggesting health-related consequences for species-specific host-microbiome interactions across hominids.

## Introduction

The microbiome of the primate gastrointestinal tract plays an important role in host physiology and health. Extreme variation in the gut microbiome has been observed between healthy human individuals; this variation is even more pronounced between different species of great apes (Human Microbiome Project Consortium, 2012; Nishida & Ochman, 2019). Microbiome composition is strongly correlated with the species of the host, a pattern known as co-diversification. Within hominids and other nonhuman primates, co-diversification between host and microbial symbionts has led to overall microbiome composition clustering along the expected phylogenetic relationships of the host species, including bacterial, archeal and eukaryotic groups within the gut microbiome (Amato, et al., 2019; Mann et al., 2019; Moeller et al., 2012; Ochman et al., 2010; Raymann et al., 2017). However, reports show that these phylogenetic constraints are flexible, depending on diet and subsistence strategy (Gomez et al., 2019). For example, compared with industrialized human groups, small scale rural or agricultural human populations share a greater number of gut microbiome traits with wild nonhuman primates (Amato, et al., 2019; Gomez et al., 2019).

Different hominid species harbor many of the same bacterial phyla in the gastrointestinal tract, but in varying abundances. For example, both the human and chimpanzee guts are primarily colonized by Bacteroidetes and Firmicutes, but the chimpanzee gut also harbors higher abundances of microbial phyla that are relatively rare in humans, including Actinobacteria, Euryarcheaota, Tenericutes, and Verrucomicrobia (Nishida & Ochman, 2019; Ochman et al., 2010). Gorillas, besides also displaying presence of these rare taxa, harbor greater abundances of Chloroflexi, Tenericutes, and Fibrobacteres (Gomez et al., 2015, 2016; Hicks et al., 2018). Although the orangutan microbiome has not been characterized as thoroughly, a previous report has shown that orangutan guts harbor higher diversity in archaeal lineages compared to other great apes, in addition to similar microbial phyla as gorillas and chimpanzees (Delsuc et al., 2014; Raymann et al., 2017). At lower microbial taxonomic levels, very different microbial species are present in human and chimpanzee microbiomes, resulting in greater divergence (Nishida & Ochman, 2019).

Overall gut microbiome composition is shaped by a combination of host genetics, host physiology, and environmental factors. Studies have shown that host genetic variation influences microbiome composition within humans, but has yet to be studied in other hominids (Blekhman et al., 2015; Goodrich et al., 2014). Among environmental influences, diet has a large impact on the primate gut microbiome (Gomez, et al., 2016; Hicks et al., 2018; Nagpal et al., 2018). Most non-human great ape species in the wild and in captivity subsist on a primarily plant-based diet of fruit and vegetation that is occasionally supplemented by animal protein, such as meat or insects (Tutin & Fernandez, 1993; Vogel et al., 2015; Watts et al., 2012). In contrast, human diets are usually omnivorous and highly variable depending on cultural influences, agricultural practices, geographic location, and individual dietary preferences (Lang et al., 2018; Wu et al., 2011). Other environmental factors that can influence microbiome composition between primates include variation in geography, seasonality, and other social behaviors such as grooming (Grieneisen et al., 2019; Tung et al., 2015). In addition, physiological differences between primate species, such as differences in gut morphology and digestive processes, also contribute to differences in microbiome composition (Amato, et al., 2019). Although a large effort has been made to characterize the factors that influence variation in the microbiome, it is unclear how variation in microbiome composition between great ape species can impact relevant host phenotypes.

A likely mechanism by which the microbiome can affect host physiology is through regulating the expression of host genes in interacting intestinal epithelial cells (Luca et al., 2018; Richards et al., 2016, 2019). Studies in animal models have demonstrated that gut microbiota can drive changes in host gene expression by altering epigenetic programming, such as histone modification, transcription factor binding, and methylation (Camp et al., 2014; Krautkramer et al., 2016; Pan et al., 2018; Qin et al., 2018). For example, Camp et al. found that the microbiome drives the differential expression of transcription factors enriched in accessible binding sites (Camp et al., 2014). In addition, Pan et al. found that the microbiome can alter DNA methylation in the gut epithelial cells of mice (Pan et al., 2018). Moreover, in cell culture, inter-individual variation in microbiome composition can drive differential responses in host gene expression at the intestinal level (Richards et al., 2019). However, we do not know how interspecies variation in the microbiome affects gene regulation in host cells. When considering the microbiota variation amongst great ape species and their influences on host gene expression, *in vivo* studies in experimental animal models are limited. Furthermore, *in vivo* experiments can be confounded by a multitude of factors, such as differences in diet between the animal model species and the primate species of interest, microbiota colonization history of the animal model, and inherent differences in the genetic backgrounds between the animal model and the primate species (Luca et al., 2018).

Here, we use an *in vitro* experimental system (Richards et al., 2016, 2019) to assess host gene expression changes in response to diverse gut microbiota from four great ape species: humans (*Homo sapiens*), and captive chimpanzees (*Pan troglodytes*), gorillas (*Gorilla gorilla gorilla*), and orangutans (*Pongo abelii*). This experimental design allows us to determine causal relationships between gut microbiome composition and gene expression changes in colonic epithelial cells that are induced by the microbiome while controlling for potentially confounding environmental and technical effects (Richards et al., 2016, 2019). We have leveraged this design to ascertain how host genes respond to between-species variation in microbiome composition across hominids, characterize the function of host genes that respond to microbiota from each great ape species, and identify microbial taxa and pathways that likely drive expression of specific host genes.

## Results

To assess how host genes respond to variation in the microbiome, we extracted live microbiota from 19 fecal samples from four hominid species (4 humans, 3 chimpanzees, 6 gorillas, and 3 orangutans), and treated human colonic epithelial cells (colonocytes) with the extracted microbiota using an experimental technique from a previously published method (see SI Table 1) (Richards et al., 2016, 2019). We quantified changes in gene expression in the colonocytes as a response to the primate microbiota using RNA-seq **(Fig. 1A)**. Additionally, we used 16S rRNA sequencing and shotgun metagenomics to characterize the composition of the microbiome in these samples. A principal coordinate analysis of Bray-Curtis dissimilarities confirmed that the microbiome samples cluster by primate host species of origin (**Fig 1B**; SI Fig. 1). This observation is consistent with previous findings showing that the phylogenetic relationship between primate host species is reflected in their microbiomes (Ochman et al., 2010), and that interspecies microbiome distinctions between wild apes is maintained in the captive individuals included in our study.

**Figure 1.**
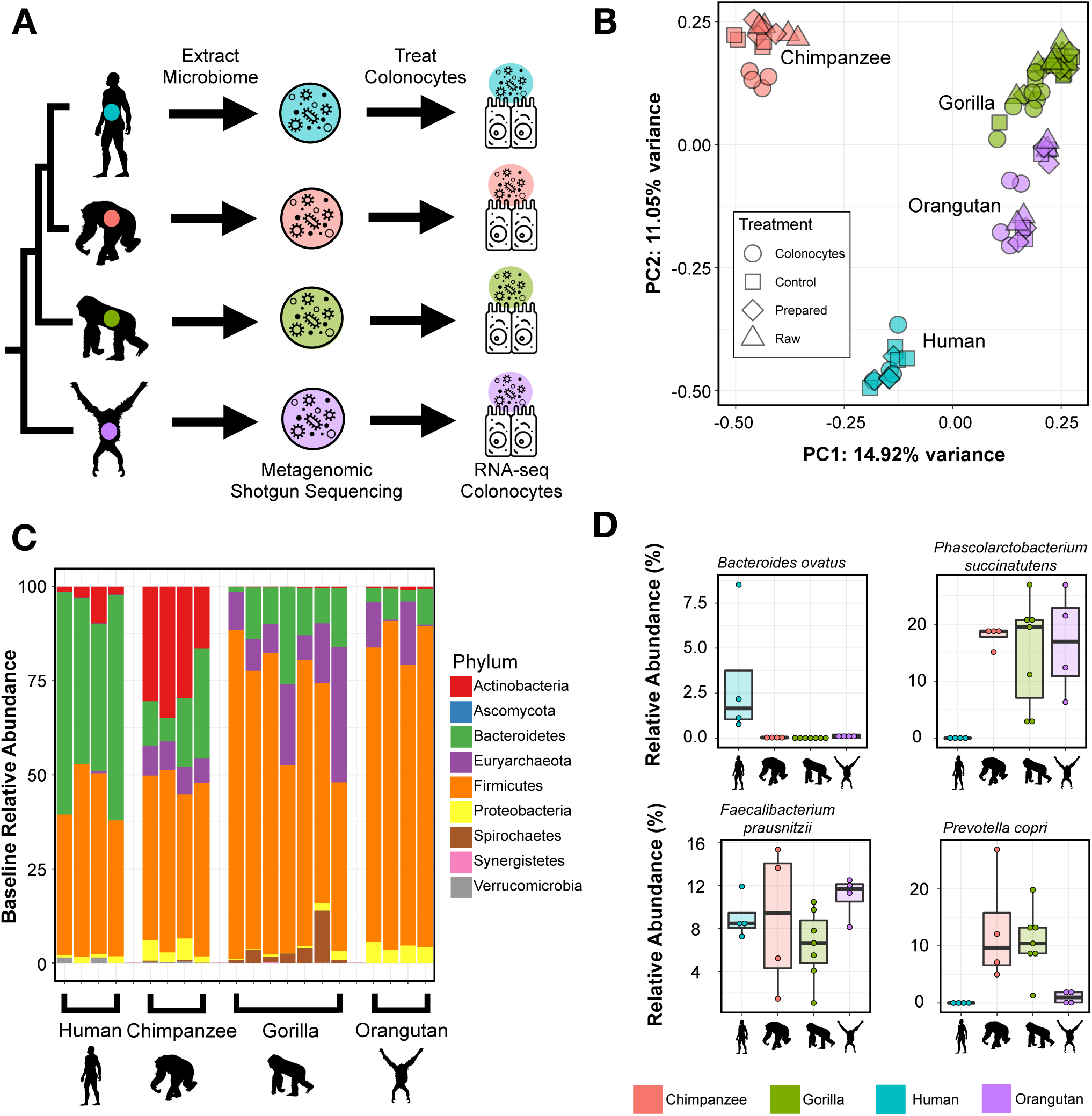
Experimental design and gut microbiome composition. **(A)** Experimental design. Live microbiomes were extracted from fecal samples from chimpanzees (n=4), gorillas (n=7), humans (n=4), and orangutans (n=4). Microbes were incubated with human colonic epithelial cells for 2 hours, after which RNA-seq was performed on the epithelial cells. Metagenomic shotgun sequencing and 16s rRNA sequencing were performed on the microbiome samples to determine microbiome composition. **(B)** Bray-Curtis dissimilarity of the all microbiome samples from all four primate species at different stages of the experiment: in raw fecal samples (raw), after extraction from fecal samples and before the experiment (prepared), after incubation with colonocytes (colonocytes), and after incubation without colonocytes (control). **(C)** Stacked barplot showing the relative abundances of microbial phyla for each hominid fecal sample prior to culturing but after extracting the microbiota for treatment. **(D)** Examples of microbial species (from shotgun metagenomics) that show various patterns of abundance across hominid species. *Bacteroides ovatus* shows a high abundance in humans relative to the other hominid species. *Phascolarctobacterium succinatutens* is highly abundant in the non-human hominid but not present in the human fecal samples. *Faecalibacterium prausnitzii* is highly abundant in all four hominid species. *Prevotella copri* is highly abundant in the chimpanzee and gorilla samples, has a lower abundance in the orangutan samples, and is not present in the human samples.

The bacterial composition of the samples confirmed clear distinctions between hominid species at the phylum level (**Fig. 1C**, SI Fig. 2), with nine of the most abundant microbial phyla showing significantly different levels between hominid species (Kruskal-Wallis test, Benjamini-Hochberg FDR <0.1, SI Table 2). The human microbiome samples have a high relative abundance of Bacteroidetes and Firmicutes, which have both been previously identified as dominant phyla in the human gut (Human Microbiome Project Consortium, 2012; Turnbaugh et al., 2007). In addition, Actinobacteria abundance is significantly different between hominid species (Kruskal-Wallis Test, Benjamini-Hochberg q-value = 0.00567; ANOVA, Benjamini-Hochberg q-value = 3.82×10), with chimpanzees showing the greatest abundance (see **Fig. 1C**). Furthermore, we identified 21 microbial species that are differentially abundant between hominid host species (SI Table 3, Kruskal Wallis, Benjamini-Hochberg FDR <0.1). Examples of several microbes that have variable abundance across species, including *Bacteroides ovatus*, which shows higher abundance in humans compared to other hominids; *Phascolarctobacterium succinatutens*, which shows lower abundance in humans compared to other hominids; and *Prevotella copri*, which has higher abundance in gorilla and orangutan, are shown in **Fig. 1D**.

**Figure 2.**
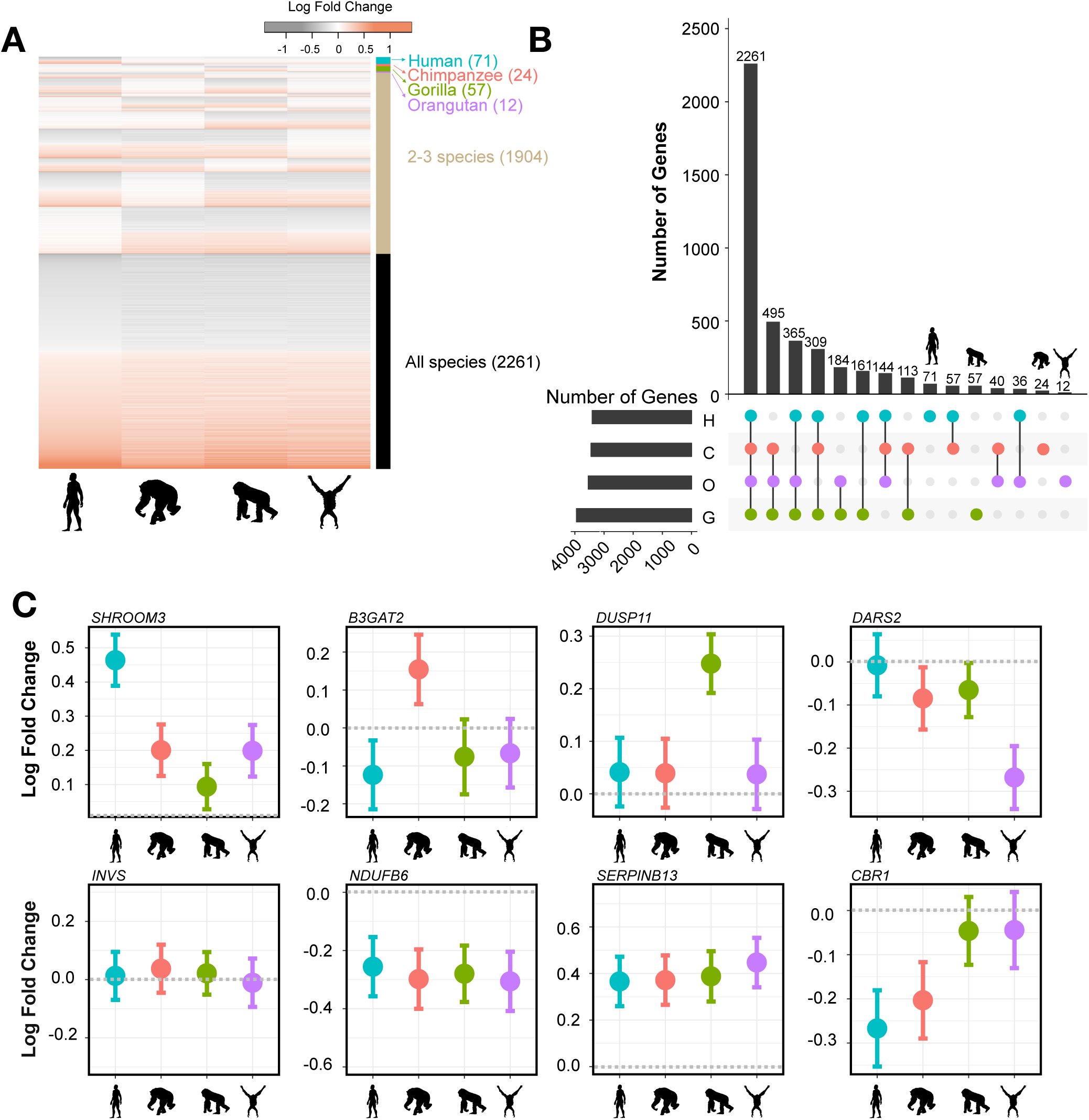
Patterns of host gene expression changes in response to hominid microbiome treatment. **(A)** Log_2_ fold change for all differentially expressed genes (rows), grouped by expression pattern. The colored bar on the right hand side indicates in response to which hominid microbiome these genes change their expression. **(B)** UpSet plot visualizing the intersections among the sets of host genes that respond to hominid microbiomes. **(C)** Examples of the expression pattern of 8 differentially expressed genes. Each panel shows the change in expression (y-axis) in response to the four hominid microbiomes (x-axis) of a single host gene, with the gene name listed at the top of each panel. Error bars indicate 1X the standard error.

To characterize the host response to the microbiome, we used likelihood ratio tests combined with a negative binomial model (DESeq2) to identify host genes that change their expression after inoculation with microbiomes from the four hominid host species (see Methods). We identified 4,329 host genes that respond to the microbiome of at least one hominid species (**Fig. 2A**, Benjamini-Hochberg FDR<0.1). The majority of differentially expressed genes (2,261 genes, 52%) respond to the microbiomes of all four hominids (see **Fig. 2A, 2B;** full dataset available in SI Table 4; see Methods). Despite this overall consistent response, we find 164 host genes that respond in a species-specific manner; namely, respond to the microbiome of one hominid species but not the other three. For example, *SHROOM3* responds to the human microbiome, but shows no response to the chimpanzee, gorilla, and orangutan microbiomes (**Fig. 2C**). Similarly, *B3GAT2, DUSP11*, and *DARS2* respond in a species-specific manner to the chimpanzee, gorilla, and orangutan microbiomes, respectively (**Fig. 2C**). We also find 394 host genes that respond to microbiomes from two hominid species; e.g., *CBR1* responds to orangutan and gorilla microbiomes (**Fig. 2C**). Likewise, 1,313 host genes respond to microbiomes from three hominid species, and 13,531 genes show no response to any of the hominid microbiomes (e.g., *INVS*; see **Fig. 2C**).

To understand how genes with a host species-specific response may interact with each other, we visualized interaction networks for differentially expressed host genes that respond to microbiomes from each hominid species (Krämer et al., 2014)(Ingenuity Pathway Analysis, http://www.ingenuity.com; **Fig. 3A, 3B**, SI Fig. 3, SI Fig. 4; see Methods). The most significant interaction network of host genes that respond only to human microbiomes is enriched with functional categories related to cancer, cell death and survival, and organismal injuries and abnormalities **(Fig. 3A**, SI Table 5). This is consistent with previous studies showing that the microbiome may influence host disease through changes in host gene regulation, but also suggests that this effect may be specific to human microbiomes (Camp et al., 2014; Krautkramer et al., 2016; Pan et al., 2018; Qin et al., 2018). By comparison, the most significant interaction network of genes that respond specifically to orangutan microbiomes is enriched for functional categories related to carbohydrate metabolism, lipid metabolism, and small molecule biochemistry **(Fig. 3B**, SI Table 6). This is consistent with the observation that orangutan diets, compared to that of gorillas or chimpanzees, could incorporate a greater proportion of ripe fruits and highly digestible/simple sugars in peak seasons (up to 100% dependence on fruit) (Remis, 1997; Taylor, 2006). In addition, previous reports point to a highly diverse archeal community in orangutans compared to other apes, which could be associated with an increased capacity to metabolize highly fermentable plant materials (Raymann et al., 2017). For functions enriched in the most significant networks for genes that respond only to gorilla microbiomes and only to chimpanzee microbiomes, see SI Table 7 and SI Table 8.

**Figure 3.**
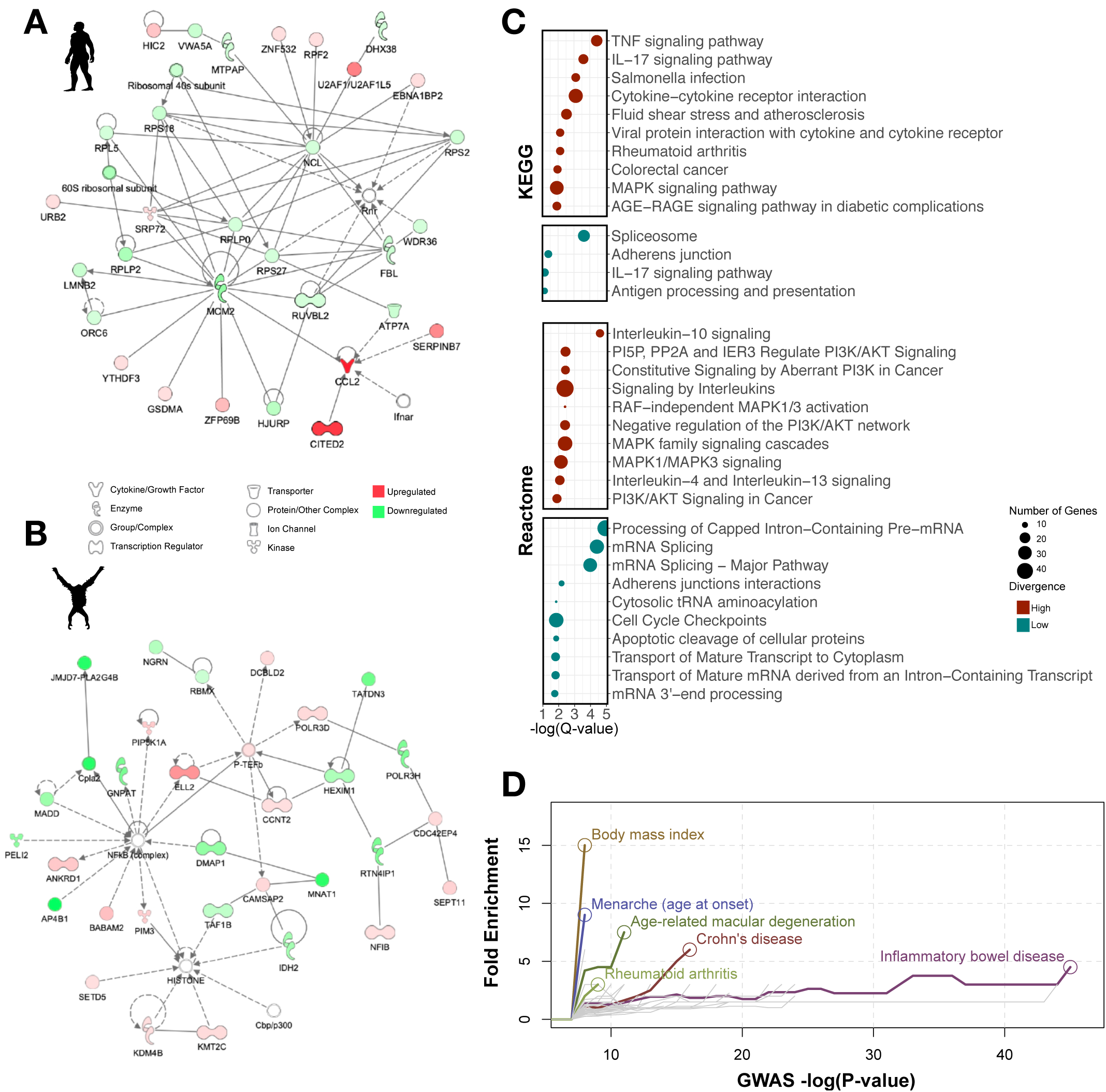
Interaction networks and functional enrichment categories for host genes responding to hominid gut microbiomes. **(A)** Interaction network showing host genes that respond only to human microbiomes, generated using Ingenuity Pathway Analysis. **(B)** Similar to (A), but including host genes that respond only to orangutan microbiomes. **(C)** Functional categories in the KEGG (top) and Reactome (bottom) databases enriched among high-divergence genes (red) and low-divergence genes (blue). X-axis indicates the statistical significance of enrichment, and the circle size corresponds to the number of genes in each category. **(D)** Complex disease enriched among genes that respond to hominid microbiomes. Fold enrichment (y-axis) is shown for a given P value threshold (x-axis) to define genes that are associated with each complex disease in the GWAS Catalog. Each colored line represents a different complex disease with an enrichment of at least three-fold, with a circle indicating the most significant P-value threshold. Diseases that did not reach significance are shown in grey lines.

**Figure 4.**
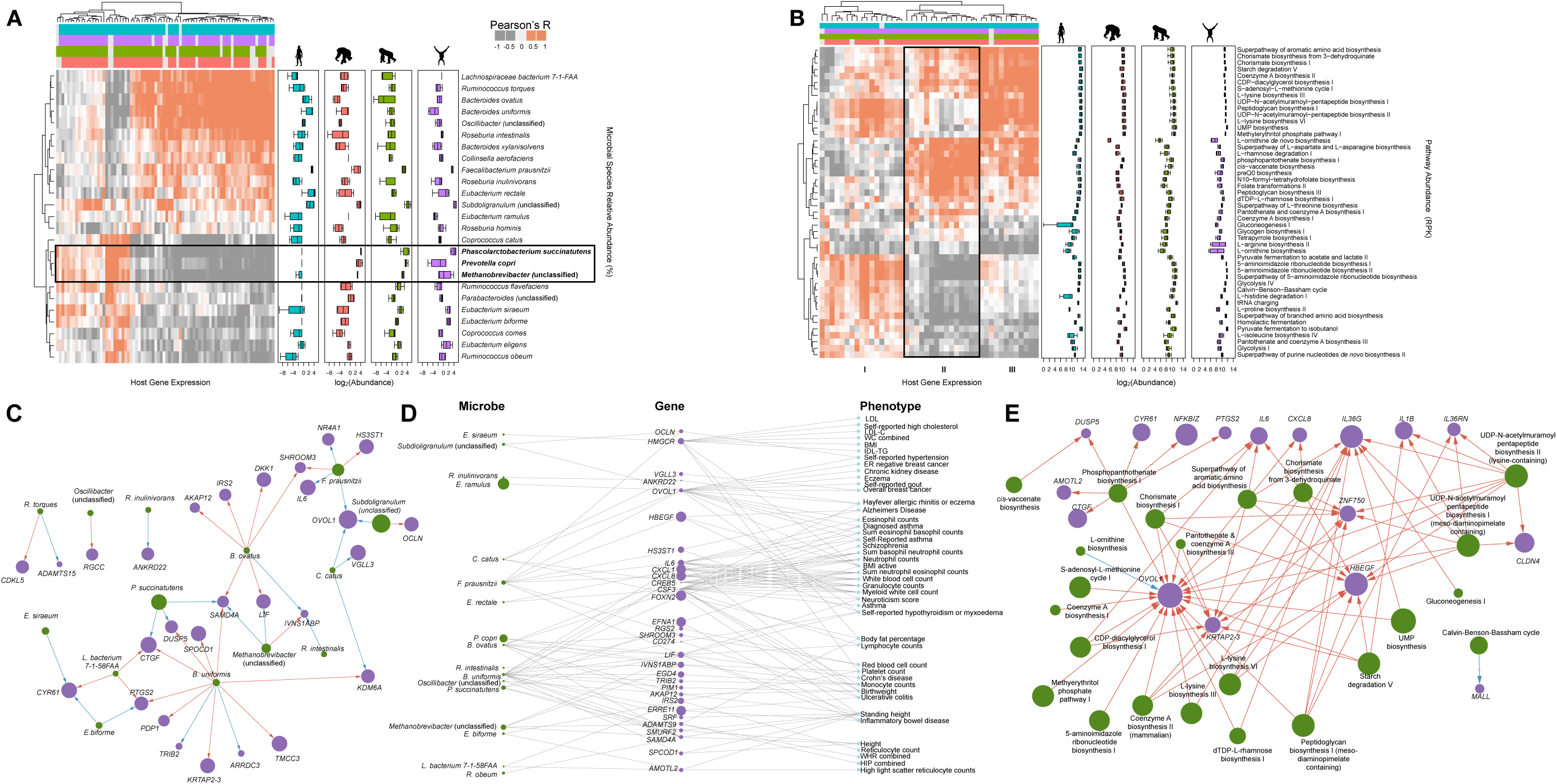
Relationship between host gene expression and specific microbiome features. **(A)** Heatmap showing correlations between microbial species (rows) and host genes (columns). Colors at the top indicate to which hominid microbiome a gene responds to. Boxplots to the right show the abundance of each microbial species in each hominid microbiome (microbial abundance transformed by log_2_). **(B)** Similar to (A), but showing microbial pathways instead of species. **(C)** Network visualization of microbial species (green) and high-divergence host genes (purple) that respond to each species connected with an arrow. Node size of microbial species and host genes corresponds to species abundance and log_2_ fold change of the differential expression, respectively. Arrow colors indicate whether a microbial species increases (blue) or decreases (red) the expression of the connected host gene. **(D)** Three tier network showing microbial species (left column), the host genes they each regulate (middle column), and TWAS phenotypes these genes are associated with (right column). Microbial species and host gene node size indicates microbial abundance and differential expression, respectively, correlated with high-divergence genes and TWAS phenotypes. **(E)** Similar to (C), but showing microbial pathways instead of species.

To further characterize the biological functions represented by host genes that respond to variation in hominid microbiomes, we categorized differentially expressed genes into two groups: low-divergence genes, which show a similar magnitude and direction of response to the four hominid microbiomes, and high-divergence genes, which show a highly variable response to the four hominid microbiomes (following the approach of Hagai and colleagues (Hagai et al., 2018); see Methods and SI Table 9). We find that low-divergence genes, namely, differentially expressed genes that show a similar response to microbiomes from all four primate species, tend to be enriched for functions related to basic cell processes, such as RNA processing, cell cycle, and RNA metabolic processing (**Fig. 3C**, Benjamini-Hochberg FDR<0.1, SI Tables 10-15). This suggests that these genes are likely involved in basic host responses to bacterial cells, rather than response to specific microbial features. Interestingly, high-divergence genes, namely, genes that respond differently to the microbiomes from the four primate host species, tend to be enriched for categories related to disease, inflammation, and cancer (**Fig. 3C**, SI Fig. 5, SI Fig. 6). Of note, colorectal cancer, rheumatoid arthritis, and *Salmonella* infection functional categories are enriched among high-divergence genes, and have all been associated with gut microbiome composition in previous studies (Dahmus et al., 2018; Ferreira et al., 2011; Scher & Abramson, 2011). Moreover, when considering host genes that have been previously associated with complex human traits through genome wide association studies (GWAS) using data in the GWAS Catalog (Buniello et al., 2019), we find that high-divergence genes are enriched with traits and diseases that have also been linked to the microbiome, such as Crohn’s Disease, Inflammatory Bowel Disease, and body mass index (**Fig. 3D**; see Methods). This might indicate that these complex disease phenotypes may be modulated by differences in composition of the gut microbial community through the regulation of these key host genes.

Next, we sought to identify genes whose response is directly correlated with the abundance of specific microbial taxa. To do so, we used mixed linear models that integrated host response transcriptomic data (via RNA-seq) and microbial species abundance information data (via shotgun metagenomics; see Methods). We identified 25 microbial species that drive the expression of 80 differentially expressed host genes across the four hominids (**Fig. 4A** and SI Table 16, 162 host gene-microbial taxon pairs in total, Benjamini-Hochberg FDR<0.05). A heatmap of the interactions reveals two roughly defined major clusters, one of which includes a subcluster of host genes that are downregulated by microbial taxa that are rare or absent in humans but present in the other hominids, such as *Prevotella copri, Methanobrevibacter* (unclassified), and *Phascolarctobacterium succinatutens* (highlighted in **Fig. 4A**; also see **Fig. 1D** for *P*. *copri* and *P*. *succinatutens* abundances across hominids). Genes that are downregulated in the presence of these microbial species are significantly enriched for several immune-related pathways, such as cytokine activity, IL-7 signaling, malaria, Legionellosis, and TNF signaling (SI Tables 17-20). Using a similar method, we identified 89 microbial pathways that drive the expression of 310 unique host genes for a total of 2061 significant microbial pathway-host gene interactions (Benjamini-Hochberg FDR<0.05). For simplicity, we focused on the top 48 microbial pathways that drive the expression of the top 44 unique host genes (**Fig. 4B** and SI Table 21), with a total of 216 microbial pathway-host genes pairs (examples of specific interactions can be found in SI Fig. 7). Clustering of this interaction data revealed three main clusters (I, II, III), with genes in cluster II associated with pathways that are more abundant in humans compared to other hominid microbiomes. These host genes are enriched in functional categories related to inflammation and infectious disease, including Legionellosis, malaria, and pertussis, and overlap with genes found in the cluster described above in the species-level analysis (SI Table 22).

To investigate specific host gene-microbe interactions, we considered the network of all high-divergence host genes for which expression is driven by microbial species (28 host genes and 14 microbial species; Benjamini-Hochberg FDR<0.01). We find that certain microbial taxa are represented in highly connected nodes and likely control the regulation of several high-divergence host genes (**Fig. 4C**). For example, two *Bacteroides* species, *B*. *ovatus* and *B*. *uniformis*, drive the expression of several host genes, including *LIF* and *DUSP5* respectively, both of which have been previously associated with inflammation (Habibian et al., 2017; Yue et al., 2015). *Bacteroides* is a highly abundant microbial genus in the human gut and is known to have mixed effects on human health (Wexler, 2007). Notably, *B*. *ovatus* is highly abundant in the human microbiome samples, but is at low abundances in the orangutan gut microbiomes and entirely absent in the chimpanzee and gorilla microbiomes (**Fig. 1D**, FDR<0.1).

To explore the possible phenotypic consequences of host genes for which expression is driven by certain microbial species, we integrated gene-trait associations identified through transcription-wide association study (TWAS). TWAS identifies associations between gene expression and complex traits by considering genetically predicted gene expression from expression quantitative trait locus (eQTL) studies and SNP-trait associations from GWAS. We considered genes implicated in 114 complex traits through Probabilistic TWAS (Zhang et al., 2019), and found that expression of 44 out of 57 high-divergence host genes is associated with 43 complex phenotypes (**Fig 4D**). These include diseases and phenotypes previously linked to the gut microbiome, including Crohn’s disease, inflammatory bowel disease, ulcerative colitis, body mass index, body fat percentage, and schizophrenia (**Fig. 4D**, SI Table 23). We found several microbial taxa that have higher abundance in the non-human microbiomes, including *P*. *copri*, and *P*. *succinatutens*, which have previously been hypothesized to have protective effects, downregulate the expression of host gene *LIF*, which has been linked to ulcerative colitis, inflammatory bowel disease, and Crohn’s disease in our TWAS analysis (Fig. 4C, Fig.4D) (De Vadder et al., 2016; Morgan et al., 2012). These results are consistent with findings from the enrichment analysis reported in **Fig. 3C** and **Fig. 3D**, where we found phenotypes related to inflammation were driven by high-divergence genes. Furthermore, we find that *Eubacterium rectale* and *B*. *ovatus*, microbes that have higher abundance in humans and that have been previously associated with inflammatory bowel disease (Noor et al., 2010; Zhang et al., 2017), upregulate the expression of *CSF3*, which has been reported as upregulated in ulcerative colitis patients (de Lange & Barrett, 2015; Hotte et al., 2012)

To investigate specific host gene-pathway interactions, we constructed a network of the most significant interactions between microbial pathways and high-divergence host genes as described above (**Fig 4E**; see Methods). We find that nine of these 17 host genes, including *DUSP5, CYR61, NFKBIZ, PTGS2, IL6, CXCL8, IL36G, IL1B*, and *IL36RN* (all displayed at the top layer in **Fig. 4E**) have been implicated in immune function or inflammation (Cox et al., 2004; Emre & Imhof, 2014; Gales et al., 2013; Habibian et al., 2017; Hörber et al., 2016; Müller et al., 2018; Onoufriadis et al., 2011; Ren & Torres, 2009; Rincon, 2012; Wang et al., 2017). We found that these genes are associated with several microbial pathways, including Phosphopanthothenate biosynthesis I, chorismate biosynthesis, UDP-N-acetylmuramoyl pentapeptide biosynthesis II (lysine-containing), and UDP-N-acetylmuramoyl pentapeptide biosynthesis I (meso-diaminopimelate containing).

## Discussion

Interactions between hominid hosts and their microbiomes have been an underexplored area of research, and the complexity of the host-microbiome relationship makes identifying the specific microbial features that causally impact the host phenotype inherently challenging. Here, we use an *in vitro* model to assess how gut microbiomes from different host species impact gene regulation, which is a likely mechanism for microbes to drive changes in host phenotype and health. Inoculating host colonic epithelial cells with live gut microbiome communities from four great ape species, we find that most host genes are regulated similarly by microbiomes from all four hominid microbiomes. However, some host genes are regulated only by microbiomes from a single hominid; these genes are enriched with immunity functions and are involved in the development of inflammatory bowel disease.

Chimpanzees, gorillas, and orangutans are our closest extant relatives, making these species an important study system for understanding human evolution as well as the genetic and environmental etiology of human-specific diseases. Distinct physiological, cognitive, and behavioral differences between primate species are hypothesized to be the result of changes in host gene regulation (Britten & Davidson, 1971; Enard et al., 2002; Gilad et al., 2006; King & Wilson, 1975). Indeed, studies have identified genes showing a species-specific expression pattern, and genes for which regulation likely evolves under natural selection (Blekhman et al., 2008; Brawand et al., 2011). Here, we show that microbiomes of different hominid species elicit different gene expression responses in the same type of intestinal epithelial cells (human colonocytes). Although we show that most host genes respond to microbiomes from different hominids in a similar manner, we also identified genes that exhibit a species-specific response. Thus, it may be tempting to hypothesize that some of the species-specific differences in gene expression observed previously are driven by interactions with the gut microbiome. These species-specific microbiome-regulated host genes might facilitate host-specific adaptations to physiological or dietary constraints; for example, our analysis indicates that genes with a response to only orangutan microbiomes are enriched for carbohydrate metabolism, lipid metabolism, and small molecule biochemistry, which suggests that the interaction of the orangutan microbiome and colonic epithelial cells may aid in digestion of specific macronutrients, especially those associated with diets rich in high-energy, highly digestible plant sources (e.g. ripe fruit).

In addition to environmental adaptations, species-specific responses to the microbiota may indicate tightly controlled symbiotic relationships that may result in disease phenotypes when altered. We find that high-divergence genes – namely, genes that respond discordantly to microbiomes from different hominid species – are enriched for traits associated with disease, such as inflammation and aberrant apoptosis. This suggests that genes with a response highly sensitive to the variation across hominid microbiomes may possibly play a role in host disease traits. These genes are also significantly associated with relevant disease traits in the GWAS catalog and in our TWAS analysis, including Crohn’s Disease (CD) and Inflammatory Bowel Disease (IBD). Significant distinctions exist in gut microbiome composition and diversity across apes with marked differences in subsistence strategies: for instance, industrialized human societies and primates in captivity have lower gut microbiome diversity and show higher incidences of noncommunicable diseases than small-scale human populations and wild non-human primates, respectively (Clayton et al., 2016; Gomez, et al., 2016). Thus, one hypothesis is that these unique features of the microbiome are causal for the development of diseases common in humans living in industrialized areas, but not in non-industrialized human populations or in non-human wild primates, such as IBD. Our results are consistent with this hypothesis, and further suggest that a mechanism by which the microbiome can affect disease risk is through regulating the expression of host genes in interacting colonic epithelial cells. For example, we found that several microbes that have lower abundance in humans compared to the other hominids, including *P*. *copri* and *P*. *succinatutens*, downregulate the expression of the gene *LIF*, which has been associated with IBD (SI Fig. 8). This suggests that these microbes may confer a protective effect through regulation of host genes, and their absence in humans is possibly detrimental. Conversely, we found that microbes that have higher abundance in humans compared to the other hominids, including *B*. *ovatus* and *E*. *rectale*, upregulate the expression of *CSF3*, which has been associated with inflammatory bowel disease (SI Fig. 8). This suggests that these microbes may have a human-specific pathogenic effect. Moreover, some of the genes we found to be regulated by the microbiome in a species-specific manner, such as *IL1B, IL6, IL36G, IL36RN*, and *CXCL8*, have been previously implicated in IBD (Gijsbers et al., 2004; Khor et al., 2011; Müller et al., 2018; Parisinos et al., 2018; Russell et al., 2016; Schulze et al., 2008), while others, such as *DUSP5, CYR61, NFKBIZ*, and *PTGS2*, have rules in immune response (Cox et al., 2004; Emre & Imhof, 2014; Habibian et al., 2017; Hörber et al., 2016).

Our ability to interpret these results in a comparative evolutionary context is limited by the unavailability of colonocytes from the non-human hominids in the study. The non-human hominids in the study are all captive, thus their microbiomes might not be representative of wild animals (Clayton et al., 2016); however, we find that the microbiomes used in this study still cluster by host species identity, and between-species variation in microbiome composition is preserved. Another limitation of our analysis is that the taxonomic profiling of metagenomic shotgun sequencing data relies on databases that are biased towards microbes residing in human microbiomes, and might impact our ability to detect and accurately quantify certain microbes in the non-human samples. Moreover, the *in vitro* approach used here represents a simplified version of the complex interactions occurring at the organismal level. Nevertheless, our approach allows for tightly controlled experimental conditions that can be tailored to the specific question of interest, by focusing on the relevant host cell type and microbiomes, and massively reducing confounding effects of cellular composition and the environment. Indeed, our approach allows controlling for various factors that may affect both the microbiome and host gene regulation, such as organismal-level variables (e.g., infection and hormones), host genetic variation, environmental factors (e.g., host diet), and oscillations and circadian dynamics in the microbiome and host gene expression.

In conclusion, we find that gut microbial communities from different hominids mostly elicit a conserved regulatory response in host cells, whereby most host genes respond similarly to hominid microbiomes. However, we also find that some host genes show a divergent response, and a number of host genes respond only to microbiomes from one hominid species and not the others. These genes are enriched in functional categories related to immunity and inflammation, and are over-represented in pathways involved in autoinflammatory diseases, such as IBD and Crohn’s disease. These results represent an important step towards understanding the causal relationships between variation in the gut microbiome across hominids and the regulation of intestinal epithelial cells. We hope that future studies will expand on this work using organoid culture and animal models to characterize the contribution of specific microbes to the development of disease through regulation of host genes.

## Acknowledgements

We would like to thank the primate zookeepers at the Ostrava Zoo, and especially Jana Pluhackova, for their help with chimpanzee fecal sample collection. We would also like to thank the primate zookeepers at the Como Park Zoo and Conservatory for their help with gorilla and orangutan fecal sample collection. Finally, we would like to thank Blekhman Lab members, and especially Rich Abdill, Beth Adamovicz, Laura Grieneisen, and Sambhawa Priya, for their comments and advice on the manuscript. This work is supported by NIH award R35-GM128716 (to R. B.) and NIH award R01-GM109215 (to F. L. and R. P.-R.). Partial funding was provided by the Czech-American Scientific cooperation (LH15175) supported by the Ministry of Education, Youth and Sports of the Czech Republic (to K. P.). This work was carried out, in part, by resources provided by the Minnesota Supercomputing Institute.

## Methods

### Sample acquisition and live microbiota extraction

See SI Table 1 for full details about the human and non-human primate fecal samples used in this analysis. Non-human fecal samples from gorillas and orangutans were collected from captive animals immediately after defecation. One orangutan who donated two samples was on a low dose of antibiotics for chronic colitis. Samples were collected as soon as possible (within an hour of defecation) into a 50mL conical tube containing 20mL of cryoprotectant solution consisting of a 50:50 mixture of glycerol and saline solution. The cryoprotectant was filter sterilized through a 0.22 μ m filter. Samples were shaken vigorously to distribute the cryoprotectant. Gorilla and orangutan samples were stored at −80°C within 1 hour after collection and shipped to the lab on dry ice. Chimpanzee samples were stored at −20°C within 1 hour of collection and then shipped to the U.S. lab on dry ice within one day. Human fecal samples were purchased from OpenBiome and arrived frozen on dry ice. The following briefly describes the protocol by which OpenBiome processes stool samples. The sample is collected by OpenBiome and given to a technician within 1 hour of defecation. The mass of the sample is measured and transferred to a sterile biosafety cabinet. The stool sample is put into a sterile filter bag, and a sterile filtered dilutant of 12.5% glycerol is added with a normal saline buffer (0.90% [wt/vol] NaCl in water). The sample solution is then introduced to a homogenizer blender for 60 s and aliquoted into sterile bottles. The bottles are then immediately frozen at −80°C. Any sample not fully processed within 2 hours of passage is destroyed.

To extract fecal microbiota from the non-human primate samples, inside a sterile low-oxygen cabinet we placed fecal material into a sterilized disposable standard commercial blender cup, added 20ml glycerol to reach approximately 30mL glycerol and 200mL normal saline buffer (0.90% [wt/vol] NaCl in water). Fecal material was blended until fully homogenized (about 1-2 min). Blended material was transferred to the same side of the membrane in a 330-micron filter bag and the liquid suspension of the bacterial community was collected on the other side of the filter. The resulting microbiota suspension was then mixed and aliquoted into small tubes and stored at −80°C.

The research and sample collection in this study complied with protocols approved through the University of Minnesota Institutional Animal Care and Use Committee.

### Colonocyte with hominid-derived microbiota treatment experiment

The experimental protocol used for the treatment of colonocytes with microbiota has previously been described in Richards et al., 2016 (Richards et al., 2016). Experiments were conducted using primary human colonic epithelial cells (HCoEpiC, lot: 9763), hereby called colonocytes (ScienCell Research Laboratories, Carlsbad, California, USA, 2950). The cells were cultured on plates or flasks coated with poly-l-lysine (PLL), according to the supplier’s specifications (ScienCell 0413). Colonocytes were cultured in colonic epithelial cell medium supplemented with colonic epithelial cell growth supplement and penicillin-streptomycin according to the manufacturer’s protocol (ScienCell 2951) at 37 °C with 5% CO_2_. At 24 hours before treatment, cells were changed to antibiotic-free medium and moved to an incubator at 37 °C, 5% CO_2_, and a reduced 5% O_2_.

Fecal microbiota were not thawed until the day of the experiment. Prior to treatment, the microbiota was thawed at 30 °C, and the microbial density (OD_600_) was assessed via a spectrophotometer (Bio-Rad SmartSpec 3000). Medium was removed from the colonocytes and fresh antibiotic-free medium was added to the cells, with a final microbial ratio of 10:1 microbe:colonocyte in each well. Additional wells containing only colonocytes were also cultured in the same 24-well plate for use as controls.

After 2 hours, the wells were scraped on ice, pelleted, and washed with cold phosphate-buffered saline (PBS) and then resuspended in lysis buffer (Dynabeads mRNA Direct kit, ThermoFisher Scientific, Waltham, Massachusetts, USA) and stored at −80 °C until extraction of colonocyte RNA for RNA-seq. We conducted both metagenomic shotgun sequencing and 16s rRNA sequencing on the microbiomes at four points: before preparation (raw), after preparation (prepared), cultured with colonocytes (colonocytes) and cultured without colonocytes (control). Human fecal microbiome samples were purchased as “prepared” from Openbiome and therefore were not sequenced raw.

### RNA-seq experiment and data processing

Poly-adenylated mRNA was isolated from thawed cell lysates using the Dynabeads mRNA Direct Kit (Ambion) following the manufacturer’s instructions. RNA-seq libraries were prepared using a protocol modified from the NEBNext Ultradirectional (NEB) library preparation protocol to use Barcodes from BIOO Scientific added by ligation, as described in Richards et al. (Richards et al., 2019). The libraries were then pooled and sequenced on two lanes of the Illumina Next-seq 500 in the Luca/Pique-Regi laboratory using the high output kits for 75 cycles to obtain paired-end reads. Reads were 80 bp in length. Read counts ranged between 12,632,223 and 36,747,968 reads per sample, with a mean of 18,726,038 and median of 16,993,999 reads per sample.

FastQC was used to determine quality of reads from raw data (FastQC, version 0.11.5). Trimmomatic was used to trim adapters. FastQC was again used to determine quality of reads after trimming of adapters (Trimmomatic version 0.33). Transcripts were aligned to database GRCh38 and was performed using HISAT2 (HISAT2 version 2.0.2)(Kim et al., 2019). After alignment, read counts ranged between 10,817,737 and 33,592,529 aligned reads per sample, with a mean of 17,142,585.72 and a median of 15,542,693.5 aligned reads per sample. Overall, the average alignment rate was ∼70% across samples (SI Fig. 9). The R ‘Subread’ package with the ‘featureCounts’ program was used to make the transcript abundance file (R version 3.3.3, Subread version 1.4.6).

### 16s rRNA sequencing

Sequencing on the 16s rRNA V4 region was performed at the University of Minnesota Genomics Center using the protocol described in Gohl et. al, 2016 (Gohl et al., 2016). DNA isolated from fecal samples was quantified with qPCR and the V4 region of the 16s rRNA gene was amplified using PCR with barcodes for multiplexing.

The forward indexing primer sequence is - **AATGATACGGCGACCACCGA**GATCTACAC[i5]TCGTCGGCAGCGTC and the reverse indexing primer sequence is - **CAAGCAGAAGACGGCATACGA**GAT[i7]GTCTCGTGGGCTCGG (where the bolded regions are the p5 and p7 flow cell adapters and [i5] and [i7] refer to the index sequence codes used by Illumina). The qPCR step starts with an initial denaturing step at 95 °C for 5 min followed by 35 cycles of denaturation (20s at 98 °C), annealing (15s at 66 °C) and elongation (1 min at 72 °C). After qPCR, samples are normalized to 167,000 molecules/μl. This is based on the volume of sample used for PCR1 (3ul), so 500,000 molecules is roughly 10x the target sequencing coverage. The next PCR (PCR1) step is similar to the qPCR step, except with only 25 cycles of denaturation, annealing and elongation. After the first round of amplification, PCR1 products are diluted 1:100 and 5μl of 1:100 PCR1 is used in the second PCR reaction. The next step (PCR2) is similar to the previous two PCR protocols, except with only 10 cycles of denaturation, annealing and elongation. Next, Pooled samples were denatured with NaOH, diluted to 8 pM in Illumina’s HT1 buffer, spiked with 15% PhiX, and heat denatured at 96 °C for 2 minutes immediately prior to loading. A MiSeq 600 cycle v3 kit was used to sequence the sample. The following Nextera adapter sequences for post-run trimming are also used. For read 1 – CTGTCTCTTATACACATCTCCGAGCCCACGAGACNNNNNNNNATCTCGTATGCCGTCT TCTGCTTG and for read 2 - CTGTCTCTTATACACATCTGACGCTGCCGACGANNNNNNNNGTGTAGATCTCGGTGGT CGCCGTATCATT

### Metagenomic shotgun sequencing

Metagenomic shotgun sequencing on prepared microbiota samples was performed at the University of Minnesota Genomics Center (UMGC). DNA samples were quantified using a fluorimetric PicoGreen assay. For a sample to pass QC, it needs to quantify greater than 0.2 ng/ul. If the samples pass QC they enter the TruSeq NexteraXT DNA library preparation queue. gDNA samples were converted to Illumina sequencing libraries using Illumina’s NexteraXT DNA Sample Preparation Kit (Cat. # FC-130-1005). 1 ng of gDNA is simultaneously fragmented and tagged with a unique adapter sequence. This “tagmentation” step is mediated by a transposase. The tagmented DNA is simultaneously indexed and amplified 12 PCR cycles. Final library size distribution is validated using capillary electrophoresis and quantified using fluorimetry (PicoGreen). Truseq libraries were hybridized to a NextSeq (either Single Read or Paired End). Clustering occurs on-board where the bound library molecules are clonally amplified and sequenced using Illumina’s SBS chemistry. NextSeq uses 2-color chemistry to image the clusters. Upon completion of read 1, a 7 base paired index read is performed in the case of single indexed libraries. If dual indexing was used during library preparation, 2 separate 8 or 10 base pair index reads are performed. Finally, clustered library fragments were re-synthesized in the reverse direction thus producing the template for paired end read 2. Base call (.bcl) files for each cycle of sequencing are generated by Illumina Real Time Analysis (RTA) software. The base call files are demultiplexed and then converted to index specific fastq files using the MiSeq Reporter software on-instrument.

### Characterizing microbiota

To identify microbial features from the metagenomic shotgun sequencing data, including taxa and pathway abundances, we used the HUMAnN2 pipeline with Metaphlan2 (HUMAnN2 v0.11.1, Metaphlan2 v0.2.6.0)(Franzosa et al., 2018; Truong et al., 2015). FastQC v0.11.7 was used to determine quality of sequencing reads before trimming. Sequencing adapters were trimmed from the raw reads using Trimmomatic (Trimmomatic v0.33) (Bolger et al., 2014). FastQC v0.11.7 was again used to determine quality of sequencing reads after trimming the sequencing adapters from the reads (SI Fig. 10). Metaphlan2 was used to assign taxonomy at all taxonomic levels to the sequencing reads in each sequencing file, and in particular to get relative abundances of microbial taxa for each sample. The HUMAnN2 pipeline utilizes bowtie v0.2.2 for read alignment (Langmead & Salzberg, 2012), DIAMOND v0.8.22 for high throughput protein alignment (Buchfink et al., 2015), MinPath (Ye & Doak, 2009) for pathway reconstruction from protein family predictions. The UniRef90 database was used for determining gene family abundances (Suzek et al., 2015). We found a total of 166 named microbial species detected in at least one sample (SI Fig. 11).

### Principal Coordinate Analysis of Samples

Using the 16s rRNA data from the fecal microbiota samples, we used the R package ‘DADA2’ (DADA2, version 1.2.2) to identify amplicon sequence variants (ASVs) from the reads(Callahan et al., 2016). DADA2 was used to filter and trim sequences from raw reads. Forward reads were trimmed to position 240 and reverse reads were trimmed to position 160. Reads were truncated at the first quality score less than or equal to 2. Reads with more than two errors were discarded after truncation. Amplicon sequences were dereplicated using the function ‘derepFastq.’ Sample composition was inferred using the ‘dada’ function. Chimeras were removed using ‘removeBimeraDenovo.’ We assigned taxonomy to the resulting ASVs using ‘assignTaxonomy.’ Using the R package ‘vegan’ (version 2.5-3), we calculated Bray-Curtis dissimilarities and plotted these as a principal coordinate analysis plot (Fig. 1B).

### Species-specific differential expression analysis

We filtered the RNA-seq counts table so that we only consider protein coding genes, reducing the number of considered genes from 60,674 to 19,715. Host genes were filtered for only protein coding genes using the R package ‘biomaRt’ with ensembl build 37. Within DESeq2 (DESeq2 version 1.14.1), RNA-seq counts were further filtered such that each gene had to be present at least once over all the samples, leaving 17860 tested genes (Love et al., 2014). DESeq2 uses a negative binomial model to model the count data while it also estimates an appropriate size factor to normalize each sample by its sequencing depth. Additionally, the overdispersion parameter governing the negative binomial distribution is estimated per each gene and using a regularization approach that can monitor outliers and adjust for the mean-variance dependency. The parameter governing the mean gene expression after adjusting to its sequencing depth is modeled as a linear combination that incorporates known batch effects (i.e., plate) and the effect of the biological variable of interest (i.e., each microbiome):

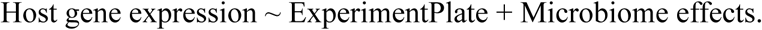

or, in mathematical terms:

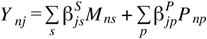

Where *Y* _*nj*_ represents the internal DEseq parameter for mean gene expression for gene *j* and experiment *n, M*_*ns*_ is the treatment indicator (control or microbiome for species *s*), and the 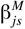 parameter is the microbiome effect for each species. To model plate as a known batch effect we use *P*_*np*_ and 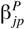 for the plate indicator variable and its effect on gene expression.

For four hominid microbiomes, 2^4^=16 effect configurations are possible (for each species combination of which parameters 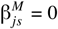), and we ran a likelihood test for each configuration *L*_*i*_ : a gene can respond to a single primate microbiome (chimp, gorilla, human, or orangutan), a gene can respond to two of the four primate microbiomes (chimp-gorilla, chimp-human, chimp-orangutan, gorilla-human, gorilla-orangutan, human-orangutan), a gene can respond to three of the four primate microbiomes (chimp-gorilla-human, chimp-human-orangutan, chimp-gorilla-orangutan, or gorilla-human-orangutan), a gene can respond to all four primate microbiomes, or a gene can show no response to any of the four primate microbiomes. The no response case is considered the base case, or null model for all the likelihood ratio tests performed.

To identify genes that respond to microbiomes from a specific primate species and to detect the total number of differentially expressed genes that respond to each of the fifteen possible non-null combinations of primate microbiomes we ran a likelihood ratio test against the base model, which assumes that the host gene shows no response: Host gene expression ∼ Experiment Plate and all the coefficients are zero. After determining across all genes and configurations which were statistically significant FDR<10%. We used the likelihood statistics *L*_*ji*_ for each gene j and configuration i to calculate the most probably configuration *P* (*H*_*ji*_|*D*) = *L*_*ji*_/Σ_*i*_*L*_*ji*_.

### Enrichment analysis

Enrichment analysis was performed using Ingenuity Pathway Analysis (IPA, QIAGEN Inc., https://www.qiagenbioinformatics.com/products/ingenuity-pathway-analysis,). We analyzed genes that show a response to microbiomes from a specific primate species. Here, we define those genes as genes that are upregulated or downregulated in response to a specific primate host species, or that show no response to microbiomes from that primate species and show a response to the other three primate host species. For example, genes that show a response only to human microbiomes will be upregulated or downregulated in response to human microbiomes, or show no response to human microbiomes and a response to chimpanzee, gorilla, and orangutan microbiomes. Genes that show a response to three species but not the fourth are also showing a species specific response to the fourth primate species.

We further validated these results using the R package ‘ClusterProfiler’ for enrichment analysis using all detected genes present in at least one sample as the background set (ClusterProfiler v3.2.14, Fig. 3C) (Yu et al., 2012). We used ENRICHR for enrichment analysis of the high and low-divergence genes and extracted the top ten response categories from the GO Biological, GO Molecular, KEGG, and Reactome databases (SI Fig. 5, SI Fig. 6)(Chen et al., 2013; Kuleshov et al., 2016).

To identify enrichment of high-divergence genes among genes that were previously found to be associated with complex human disease and traits, we used data from the GWAS catalog (Buniello et al., 2019). Since each GWAS has a different distribution of p-values and significance cutoffs, we chose to use a set of −log_10_(p-value) cutoffs in the range of 8-50 (plotted along the x axis in Fig. 3D). For a given trait, we identified the overlap between the genes significantly associated with the disease at each cutoff and high-divergence genes34, and calculated a fold enrichment (plotted along the y axis in Fig. 3D), defined as the ratio of observed/expected overlap between the two gene sets. We used a Fisher Exact Test to calculate a p-value for each cutoff, and traits for which this value was significant after FDR corrections were marked with a colored line in Fig. 3D.

K-means clustering was performed using the ‘kmeans’ function in base R (version 3.3.3) on the cluster of microbes *P*. *copri, Methanobrevibacter* and *P*. *succinatutens* for the genes in Fig. 4A. Enrichment analysis was performed using ENRICHR on the two clusters of genes. A k-means clustering analysis was also performed on the full set of microbial pathway-host gene correlations in Fig. 4B to produce three clusters of genes.

### Log fold change of genes by primate species

To calculate the fold changes for each gene for each of the four primate species, we used a similar DESeq2 model to the one described above:

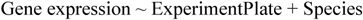

or, in mathematical terms:

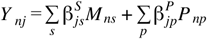

Here, Species is a vector indicating which primate species the microbiome sample originated from, and ExperimentPlate controls for the batch effect as before, but we just test the marginal effect of each species-specific parameter 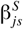 being not different than the untreated control 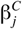. We use the contrast argument in DESeq2 to extract comparisons of each primate species against the control. Thus, this resulted in log fold change calculations for each gene as it responds to each of the four primate species’ microbiomes. These values are available in SI Tables 24-27.

### Divergence scores for differentially expressed, conserved genes

Using DESeq2, we identified genes that responded to microbiome treatment. The model to determine whether a gene responds to treatment, we used the following model:

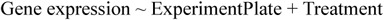

Where ExperimentPlate controls for the batch effect of the experiment, and Treatment is a binary vector indicating whether the colonocytes are treated with a microbiome or act as a control for the experiment. Mathematically:

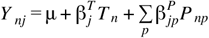

Where *Y* _*nj*_ represents the internal DEseq mean gene expression parameter for gene *j* and experiment *n* as before, *T*_*n*_ is the treatment indicator (control = 0 or microbiome = 1), and the 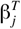 parameter is the microbiome effect. Plate effects are modeled as before. To model plate as a known batch effect we use *P*_*np*_ and 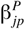 for the plate indicator variable and its effect on gene expression.

Log fold changes for each gene were calculated as described above, and then used to calculate a divergence metric for each gene. We used a similar divergence calculation as described in Hagai et al. (Hagai et al., 2018). Namely, for the genes identified as responding to treatment with microbiomes, we used the log fold changes for each species in the following equation:

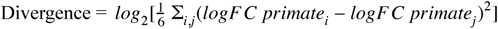

Following Hagai et al., the top 25% of genes were assigned a “high-divergence” status, and the lowest 25% of genes were assigned a “low-divergence” status. These genes were used in the enrichment analyses described below.

The rest of the genes are considered “medium divergence” genes. These genes are used in the enrichment analysis as a background set (Fig. 3A, Fig. 3C).

### Pairwise correlations between host genes and microbial species and pathways

Using the microbial species abundances calculated from the metagenomic shotgun sequencing, we ran correlation analysis between genes that are differentially expressed with respect to treatment with microbiota and abundances of microbial species. Metaphlan2 reports microbial species as a proportion of the total microbial community per sample. Microbial species were filtered such that only microbial species present in at least half of the samples and that reached a total summed relative abundance of 9% were included in the analysis, leaving 36 microbial species. We applied a center log-ratio transformation to the filtered microbial species abundance data. Microbial pathways were filtered such that the total of each pathway had to be greater than a summed threshold of 8000 reads per kilobase (RPK), leaving 95 microbial pathways to be included in the analysis. Microbial pathways were normalized using the centered log ratio transformation in a similar manner to the microbial species.

Using DESeq2, we identified which microbial species or pathways are associated with differentially expressed genes using the following model:

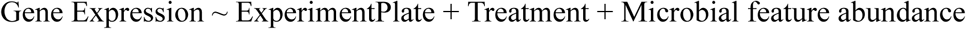

Mathematically:

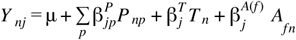

Where *Y nj* represents the internal DEseq parameter for gene expression for gene *j* and experiment *n* as before, *T*_*n*_ is the treatment indicator (control = 0 or microbiome = 1), and the 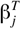 parameter is the microbiome effect. Plate effects are modeled as before. The parameter 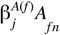 is used to model the effect of the microbiome feature (i.e., microbial species or pathway) *f* on gene expression. We statistically test effect 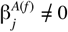 in a separate DESeq model run for each feature *f*. We used an FDR correction on the combined results from all models. The microbial species abundance is a continuous numeric value that represents that center log ratio transformed relative abundance of the microbial feature *f Afn* for each sample *n*.

### TWAS analysis

To directly investigate whether discovered effects on gene expression may contribute to complex traits, we considered PTWAS gene-trait associations for 114 traits from Zhang et al.(Zhang et al., 2019). PTWAS utilizes probabilistic eQTL annotations derived from multi-variant Bayesian fine-mapping analysis of eQTL data across 49 tissues from GTEx v8 to detect associations between gene expression levels and complex trait risk. Using the host genes that were highly correlated with a microbial species and fell into the high-divergence category (FDR<0.05), we overlapped the significant results with genes causally implicated in complex traits across all tissues by Zhang et al. (PTWAS scan, 5% FDR). We repeated the same analysis with the host genes that were highly correlated with a microbial pathway (FDR<0.01) and fell into the high-divergence category.

## Data Availability

Raw data for 16s rRNA sequencing, RNA-sequencing, and metagenomic shotgun sequencing are available on the Sequence Read Archive (SRA) under submission ID SUB7918466. For data tables used in this analysis, including tables for RNA-seq gene expression counts, metagenomic shotgun sequencing species abundances, amplicon sequence variant abundances, pathway abundances and metadata, see our Figshare project under the same title as the manuscript at https://figshare.com/account/home#/projects/87626.

