## Supplementary Figures for "Interspecies variation in hominid gut microbiota controls host gene regulation"

### Supplemental Figures

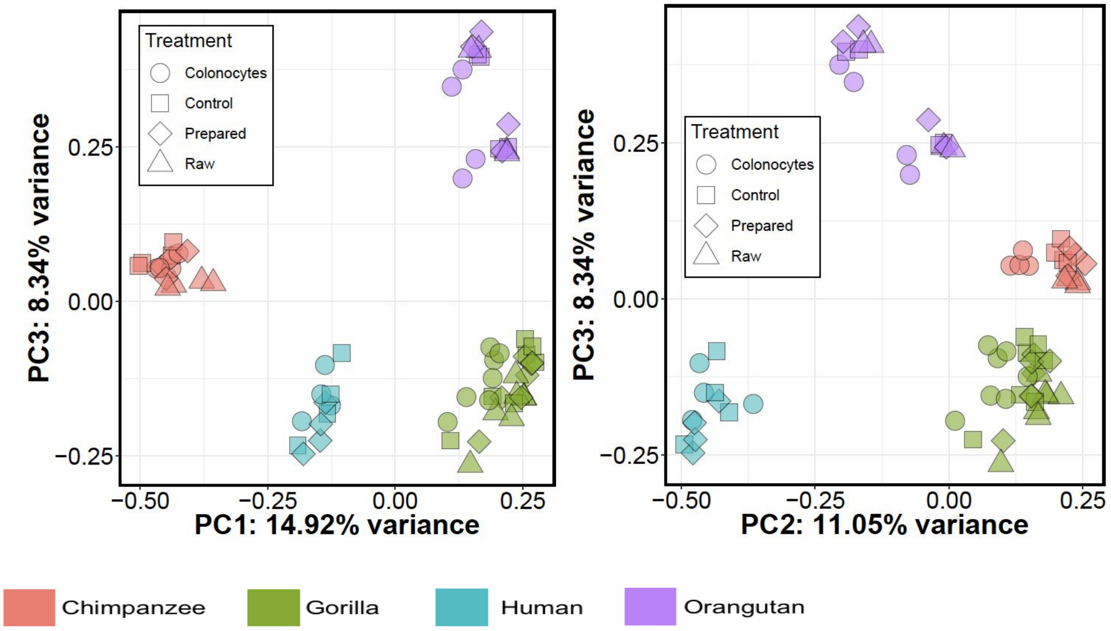


**SI Fig. 1**

PCOA plots generated using DADA2, Bray-Curtis Dissimilarity metric for primate microbiomes from all four primate species at different stages of the experiment. The left plot compares PC1 and PC3 and the right plot compares PC2 and PC3.

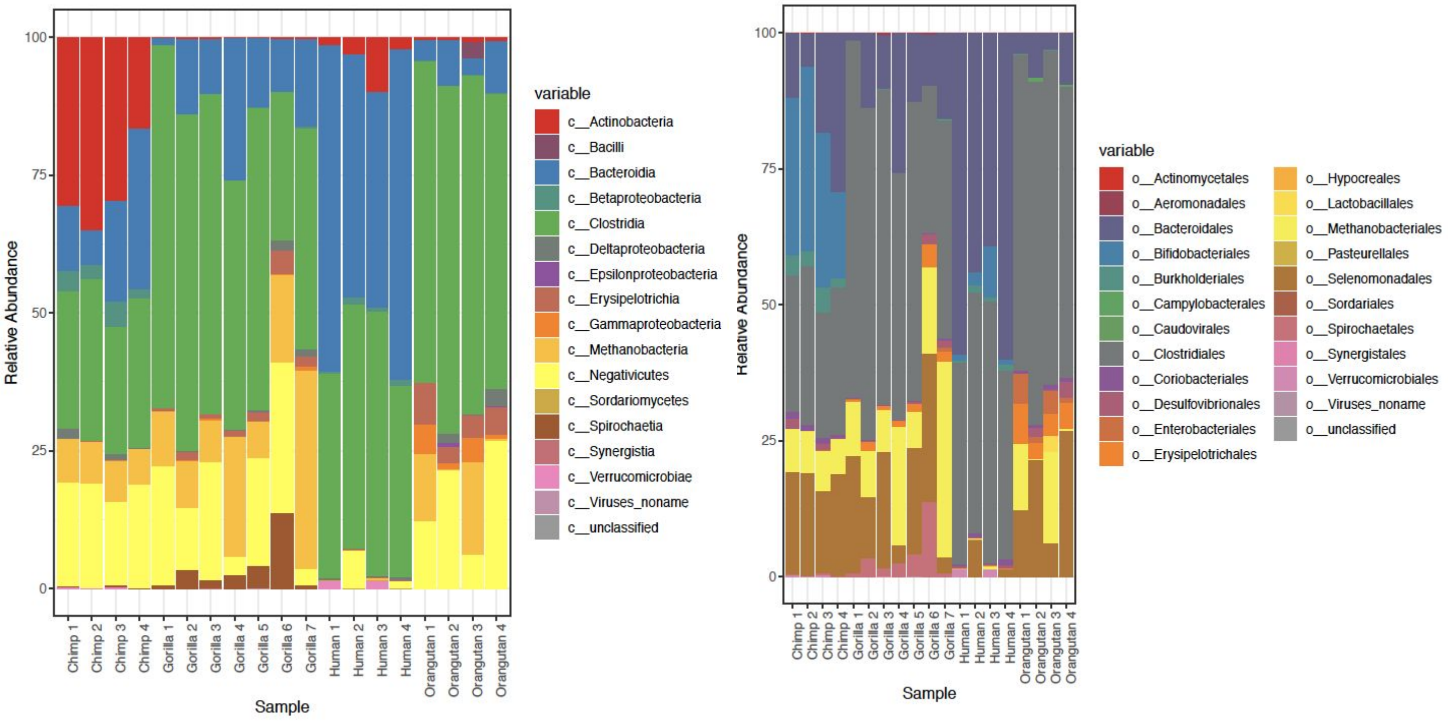


**SI Fig 2**

Barplots for the relative abundances of microbial taxa for each individual microbiome before culturing with colonocytes, at the (A) class and (B) order taxonomic levels using metagenomic shotgun analyzed with HUMAnN2.

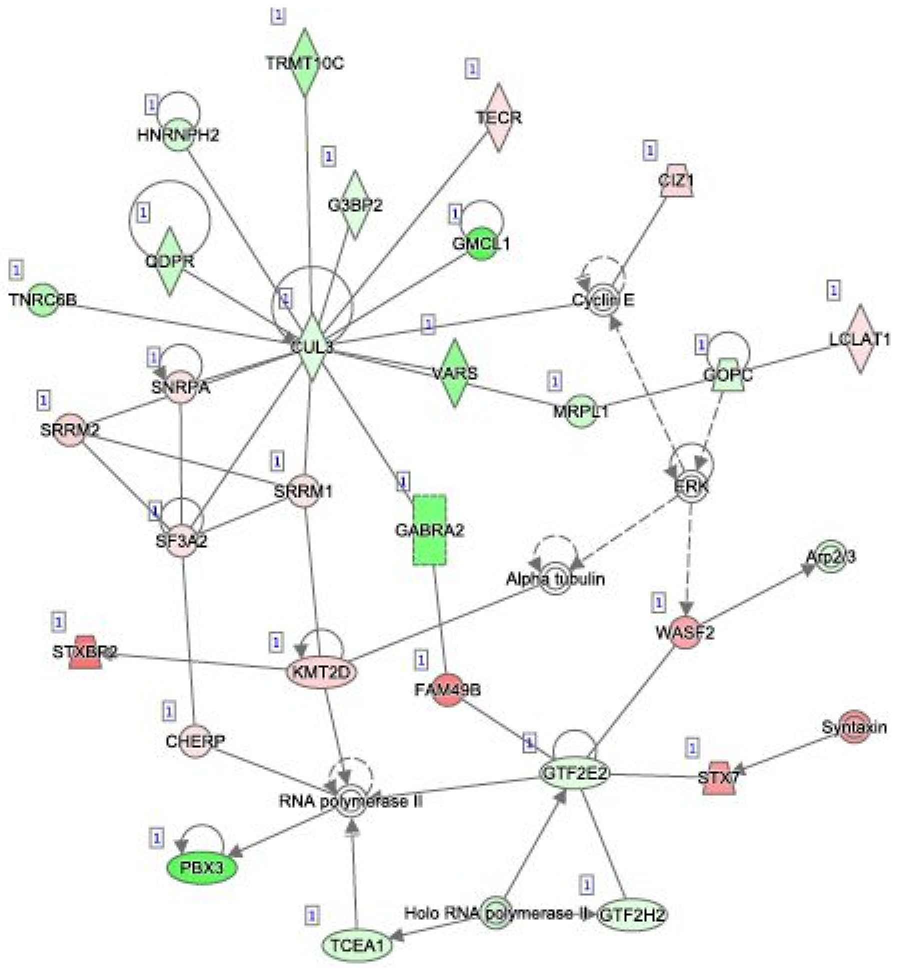


**SI Fig. 3**

Gorilla network generated in IPA which corresponds to enrichment for Cancer, Organismal Injury and Abnormalities, Skeletal and Muscular Disorders.

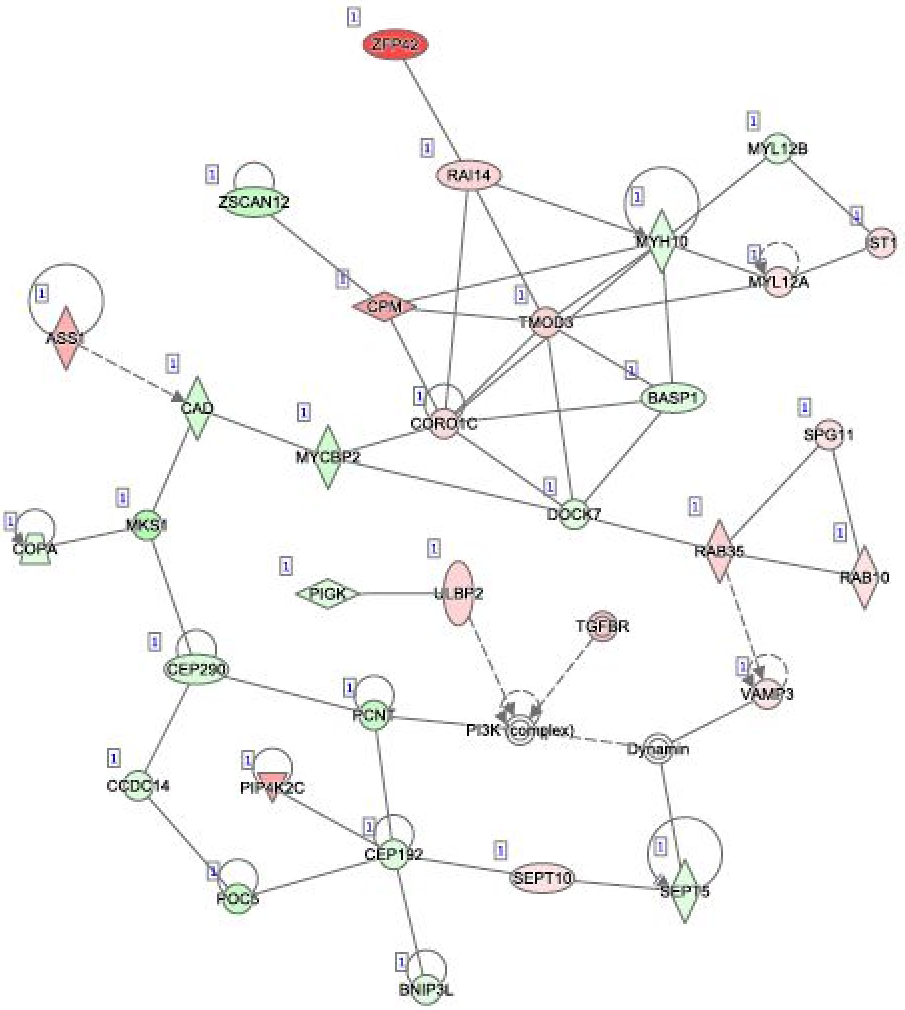


**SI Fig. 4**

Chimpanzee network generated in IPA which corresponds to enrichment for Cellular Assembly and Organization, Cellular Function and Maintenance, Developmental Disorder.

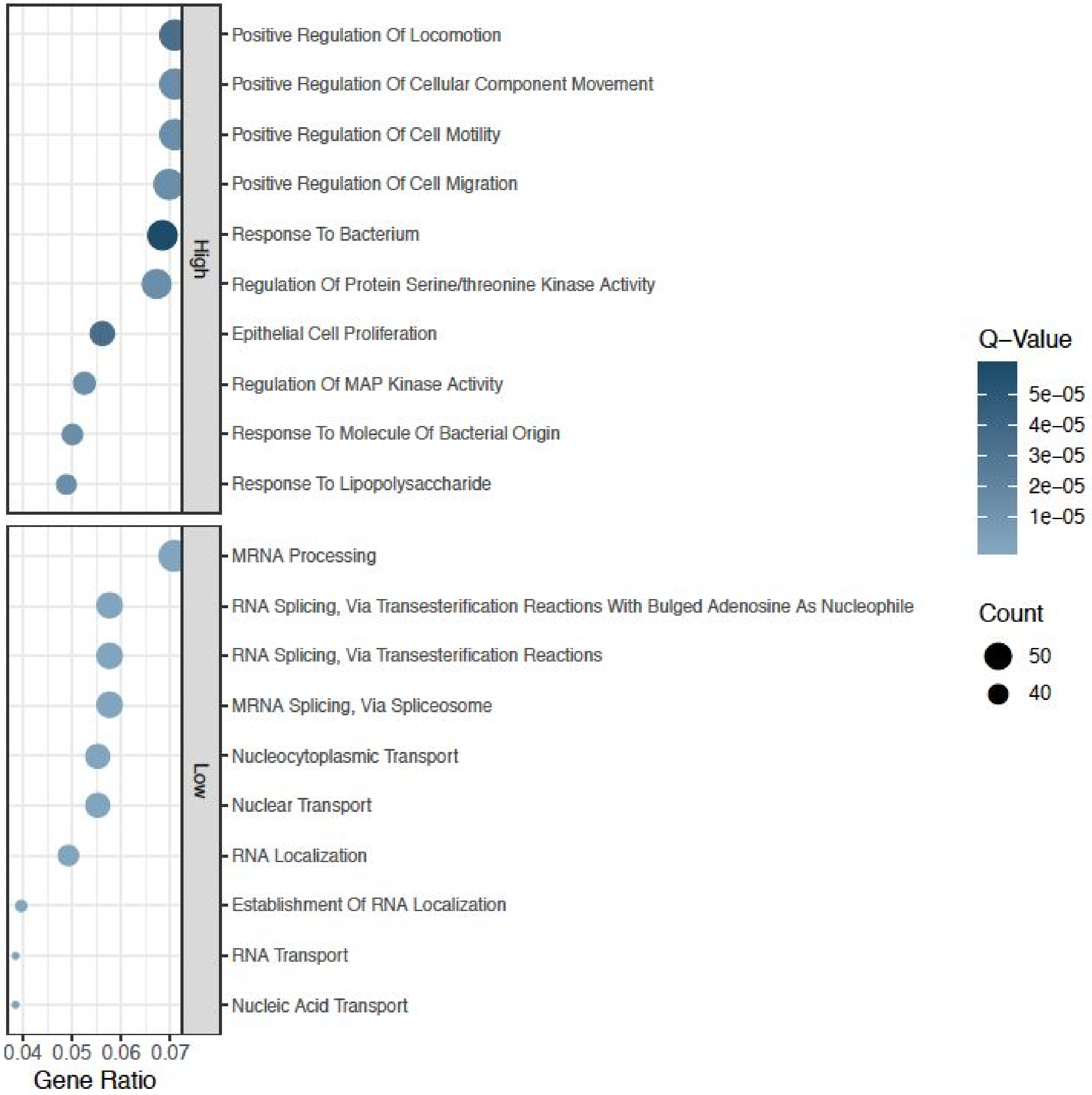


**SI Fig. 5**

Top ten Gene Ontology (GO) enrichment categories for the high-divergent genes and top ten GO enrichment categories for low-divergent genes using ClusterProfiler results.

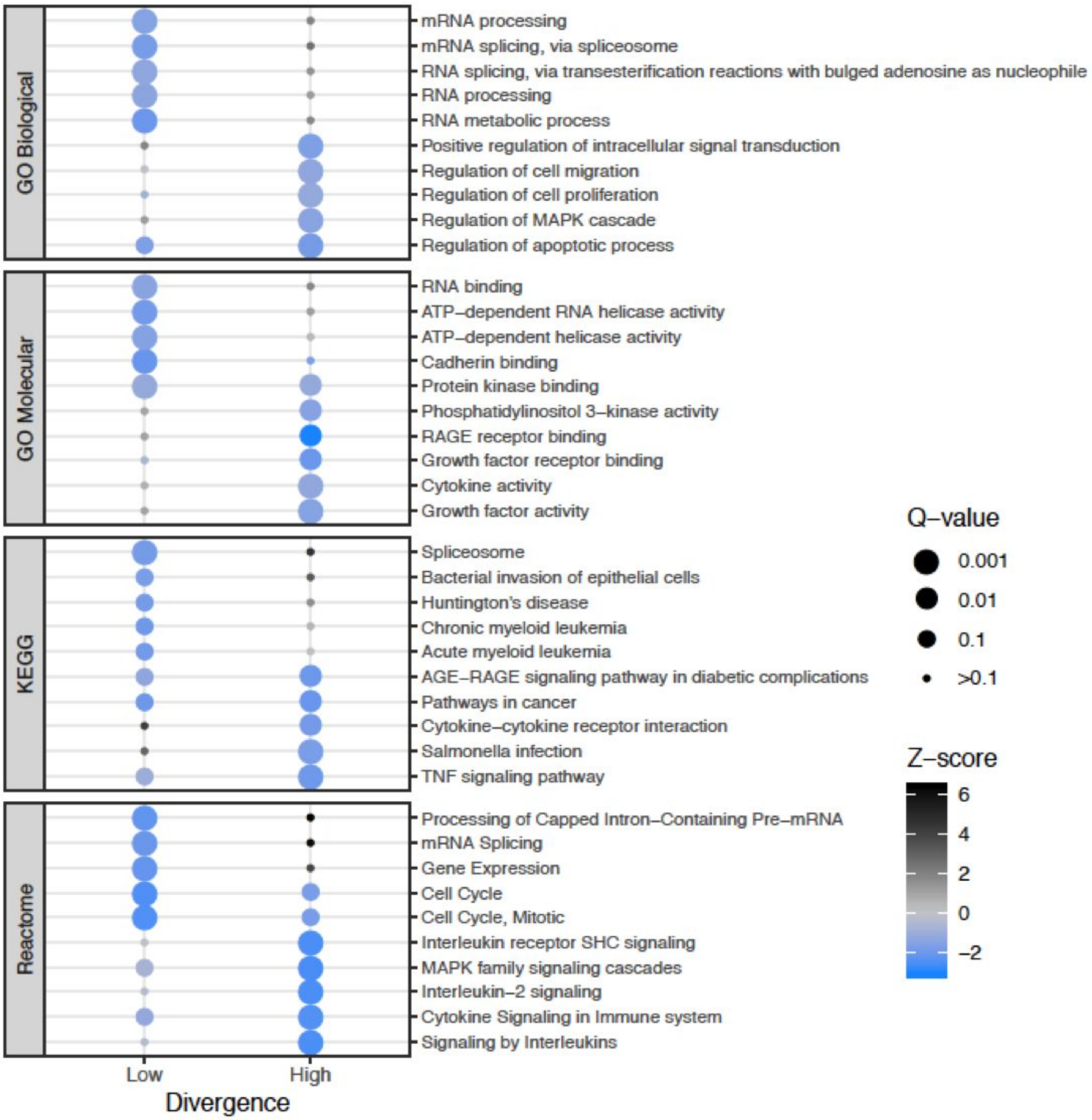


**SI Fig. 6**

Enrichment results comparing highly and lowly divergent genes using ENRICHR results for the GO Biological, GO Molecular, KEGG and Reactome databases.

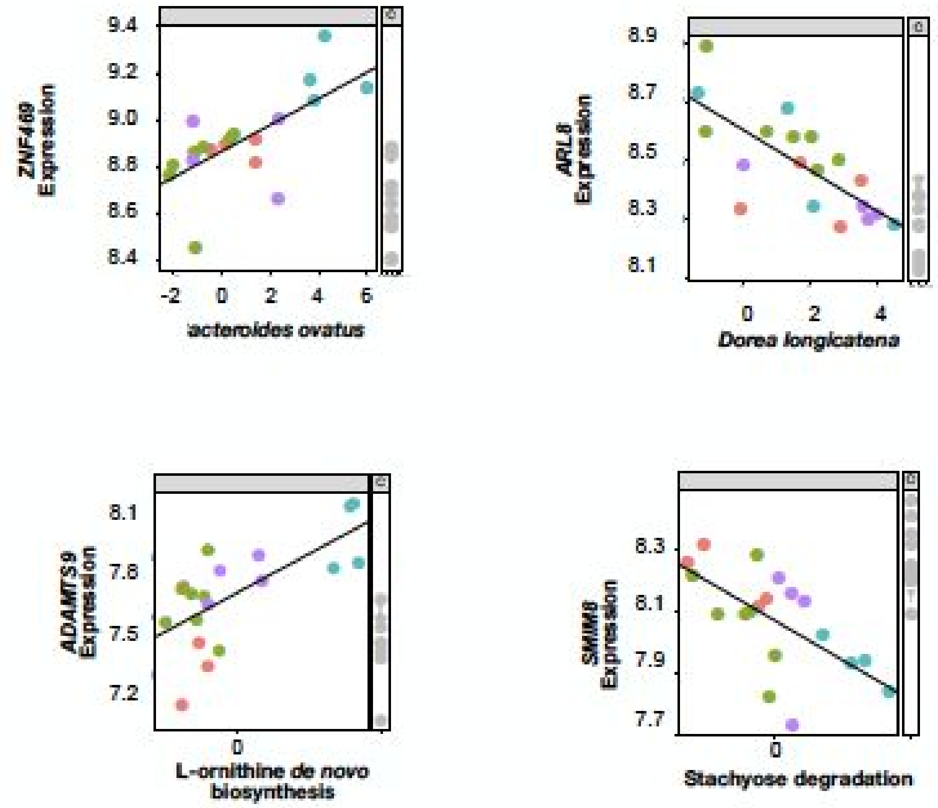


**SI Fig. 7**

Examples of microbial features plotted against host gene expression changes. The top two plots show examples of the log transformed abundance of a single microbial species plotted against the transformed log fold change of a specific gene. Each point represents a microbiota sample and is colored by the primate species of origin. The bottom two plots show examples of log transformed abundance of a single microbial pathway plotted against the transformed log2 fold change of a specific gene.

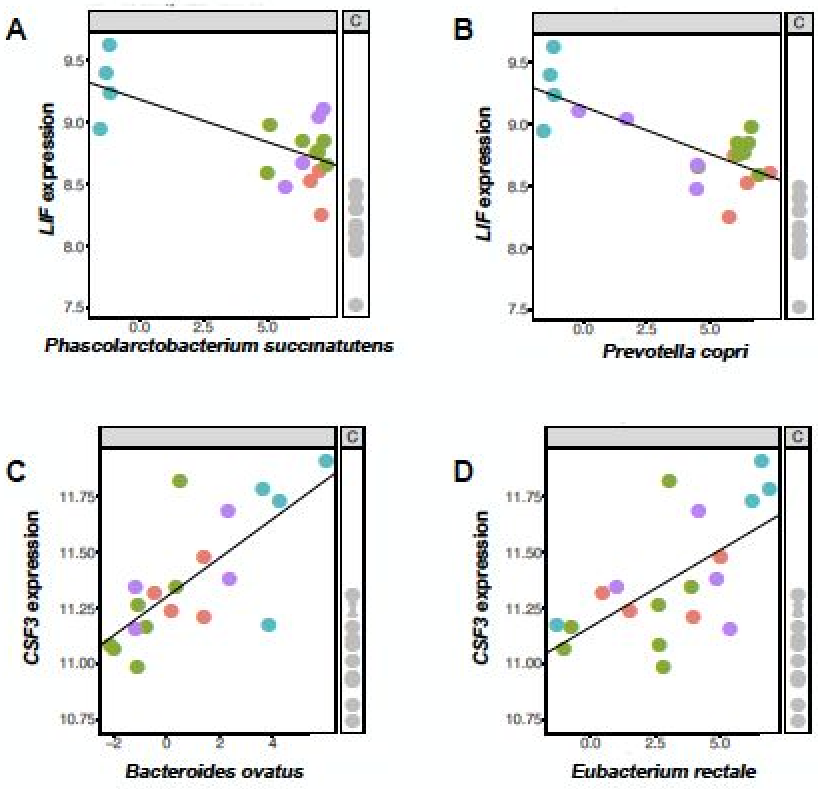


**SI Fig. 8**

Examples of microbial species and host gene interactions that are associated with inflammatory bowel disease (IBD). *Phascolarctobacterium succinatutens* and *Prevotella copri* downregulate the expression of *LIF*, while *Bacteroides ovatus* and *Eubacterium rectale* upregulate the expression of *CSF3*. Each point represents a microbiota sample and is colored by the primate species of origin.

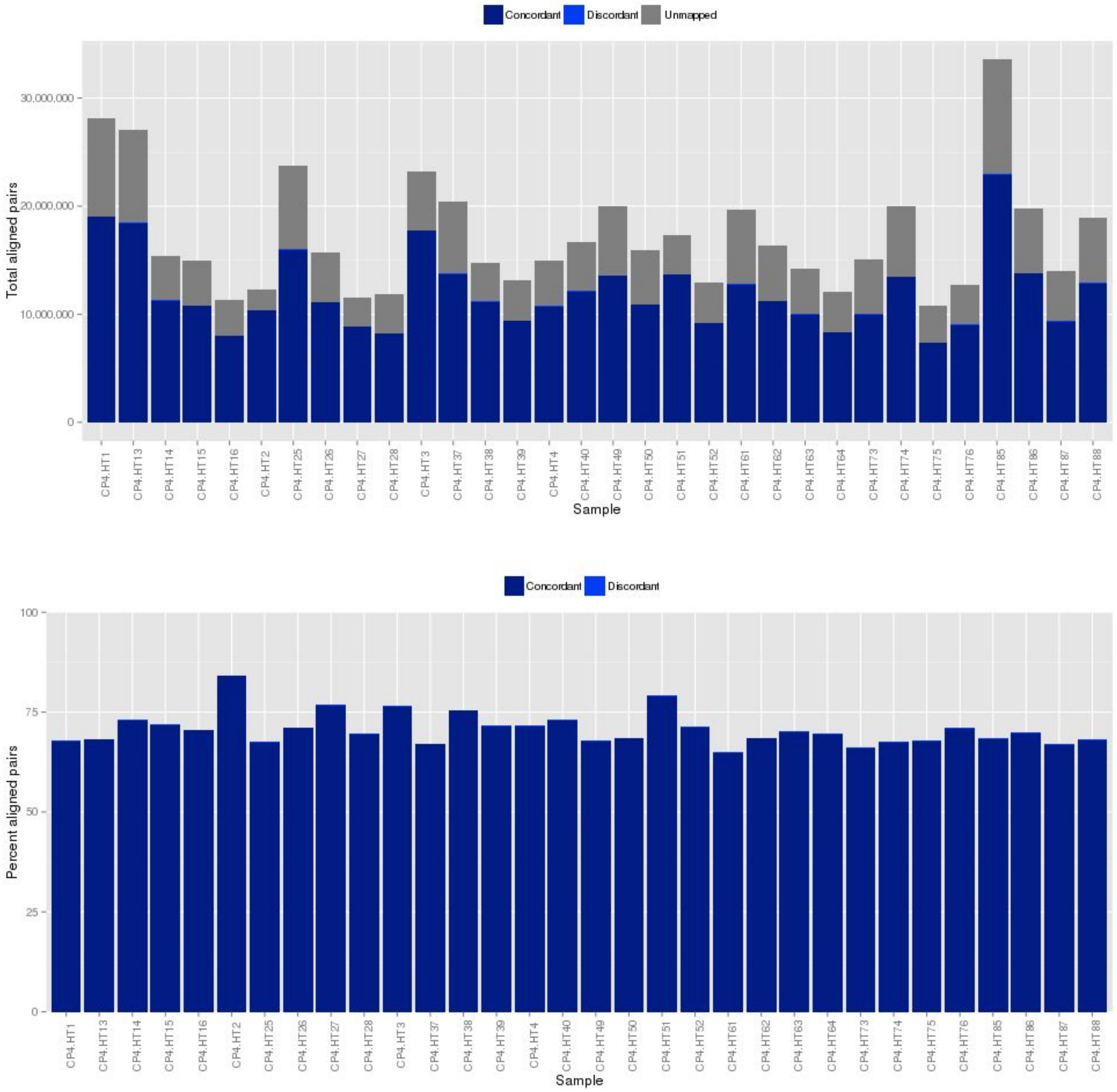


**SI Fig. 9**

Aligned Reads RNA-seq data for each colonocyte sample, cultured with or without primate microbiota. The top plot shows the total aligned pairs for each sample, and bottom shows the percent of aligned pairs for each sample.

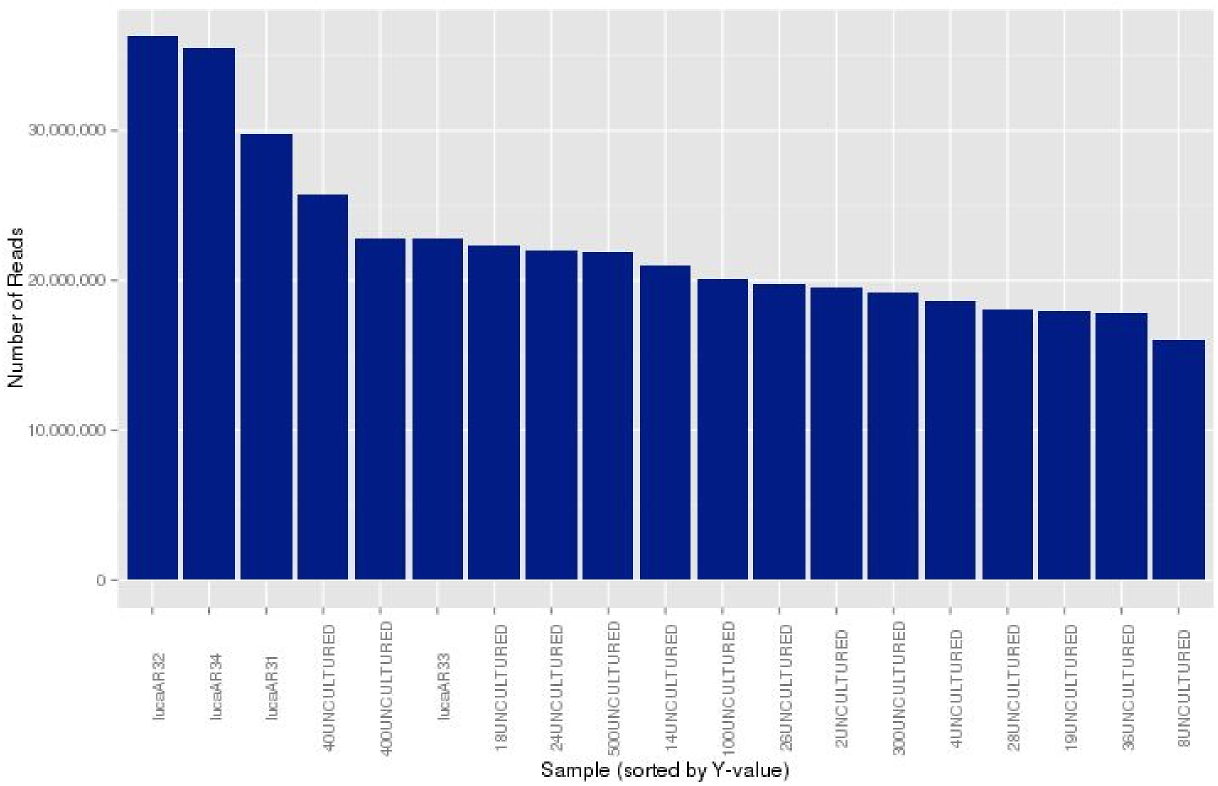


**SI Fig. 10**

Number of reads for each prepared microbiota sample based on metagenomic shotgun sequencing data.

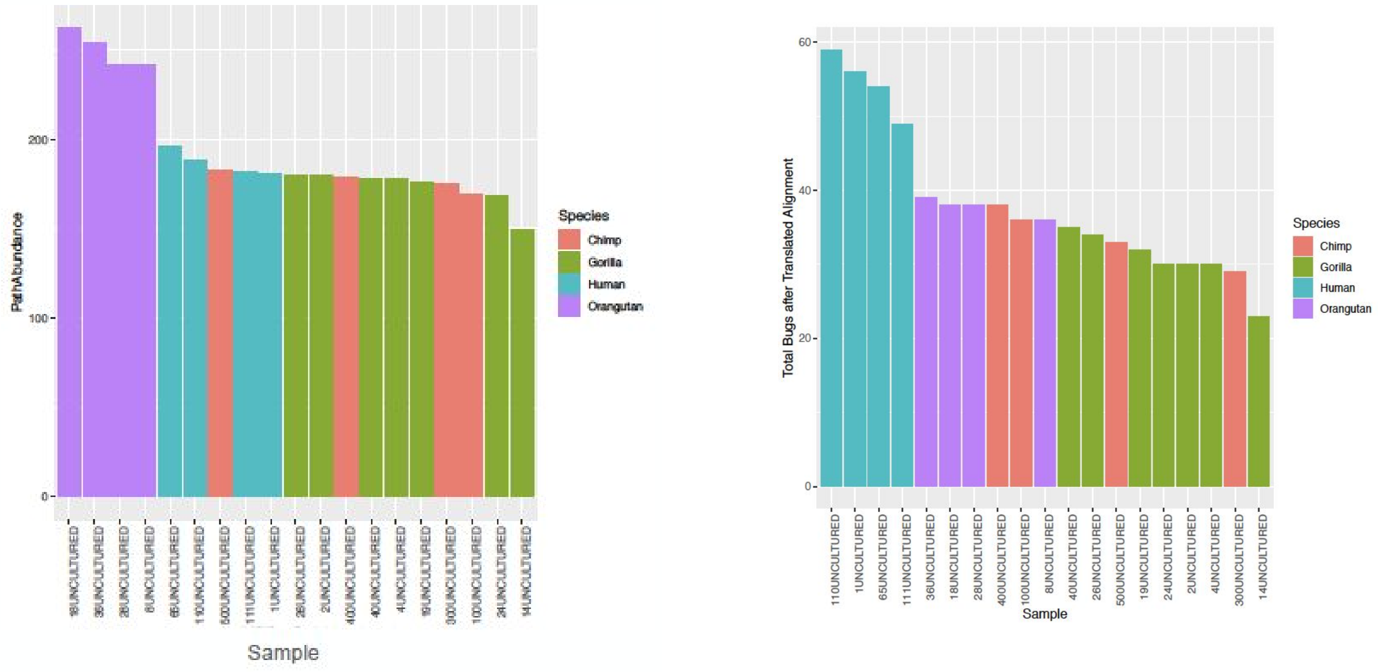


**SI Fig. 11**

From HUMAnN2 analysis to characterize each prepared primate microbiota sample. The left plot shows the number of unique microbial species after alignment for each microbiota sample, and the right plot shows the number of unique microbial pathways for each sample.
